# Subgroups of Eating Behavior Traits Independent of Obesity Defined Using Functional Connectivity and Feature Representation Learning

**DOI:** 10.1101/2022.03.03.482759

**Authors:** Hyoungshin Choi, Kyoungseob Byeon, Jong-eun Lee, Seok-Jun Hong, Bo-yong Park, Hyunjin Park

**Affiliations:** Department of Electrical and Computer Engineering, Sungkyunkwan University, Suwon, Republic of Korea; Center for Neuroscience Imaging Research, Institute for Basic Science, Suwon, Republic of Korea; Center for the Developing Brain, Child Mind Institute, New York, U.S.A; Department of Biomedical Engineering, Sungkyunkwan University, Suwon, Republic of Korea; Department of Data Science, Inha University, Incheon, Republic of Korea; School of Electronic and Electrical Engineering, Sungkyunkwan University, Suwon, Republic of Korea

**Author notes:** **Corresponding Authors:** Bo-yong Park, Ph.D. Department of Data Science, Inha University, Incheon, Republic of Korea, Hyunjin Park, Ph.D. School of Electronic and Electrical Engineering, Sungkyunkwan University, Suwon, Republic of Korea.

**Keywords:** eating behavior, subgroup, functional connectivity, autoencoder, manifold learning, representation learning, integrated gradient

## Abstract

Eating behavior is highly heterogeneous across individuals, and thus, it cannot be fully explained using only the degree of obesity. We utilized unsupervised machine learning and functional connectivity measures to explore the heterogeneity of eating behaviors. This study was conducted on 424 healthy adults. We generated low-dimensional representations of functional connectivity defined using the resting-state functional magnetic resonance imaging, and calculated latent features using the feature representation capabilities of an autoencoder by nonlinearly compressing the functional connectivity information. The clustering approaches applied to latent features identified three distinct subgroups. The subgroups exhibited different disinhibition and hunger traits; however, their body mass indices were comparable. The model interpretation technique of integrated gradients revealed that these distinctions were associated with the functional reorganization in higher-order associations and limbic networks and reward-related subcortical structures. The cognitive decoding analysis revealed that these systems are associated with reward- and emotion-related systems. We replicated our findings using an independent dataset, thereby suggesting generalizability. Our findings provide insights into the macroscopic brain organization of eating behavior-related subgroups independent of obesity.

## Introduction

Eating behavior is a key trait associated with an individual’s health [1,2]. Aberrant eating behavior can lead to a high body mass index (BMI) and cause obesity-related pathologies, such as diabetes, hypertension, and stroke [3,4]. To assess the link between eating behavior and obesity, existing studies have examined several factors that affect an individual’s eating behaviors, such as hormone activity, gene enrichment, and environmental factors [4–8]. Eating behavior is highly heterogeneous across individuals, and thus, a systematic analysis is necessary to assess individual variability.

Magnetic resonance imaging (MRI) is used to investigate brain networks associated with eating behaviors *in vivo* [9–11]. In particular, resting-state functional MRI (rs-fMRI) reflects functional alterations in the brain via temporal fluctuations in brain signals. Our previous study demonstrated associations between disinhibited eating behaviors and functional connectivity in the frontoparietal network [10]. Other studies have proved associations of eating behaviors with the brain function of the prefrontal cortex, orbitofrontal cortex, and amygdala [12–15]. These findings suggest that eating behavior is associated with the brain function. However, no clear trends were observed. Some studies have shown positive associations between disinhibited eating and brain function in the reward network [12,13], whereas others have indicated opposite patterns [14,15]. This inconsistency may be owing to the heterogeneity of eating behavior traits. Thus, the brain function differences between individuals depending on the eating behavior needs to be investigated systematically.

One approach to exploring the heterogeneity of eating behaviors is clustering, which is an unsupervised machine learning technique that defines distinct clusters with relatively homogeneous data points. Clustering techniques were widely adopted in existing neuroimaging studies to identify subgroups of healthy and diseased populations [16–20]. Some studies have classified individuals with an autism spectrum disorder into several subtypes based on cortical morphologies [21] and functional connectivity [16,17]. In addition, the clustering approach was effective in schizophrenia to assess clinical heterogeneity [19,20]. Clustering techniques are purely data-driven approaches, free from an a priori hypothesis. Thus, they can be used for identifying subgroups of a particular dataset with homogeneous characteristics. We hypothesized that clustering based on neuroimaging features may identify distinct subgroups that may exhibit different clinical or behavioral traits.

Functional connectivity is a widely adopted measure to assess co-fluctuations of the brain signals, which is defined by calculating correlations of time series between brain regions. A recent study suggested a method for characterizing functional connectivity based on manifold learning techniques [22]. These techniques produce low-dimensional representations of functional connectivity by estimating principal components based on principal component analysis or scaled eigenvectors that are based on diffusion embedding in a newly defined low-dimensional space. The generated eigenvectors exhibited smooth transitions of connectome organization along the cortical mantle, and the principal eigenvector consisted of a cortical hierarchy expanding from low-level sensory to higher-order association networks [22]. These eigenvectors have been suggested as potential imaging biomarkers in studies on healthy aging [23,24] and neurodevelopment [25–30]. In our previous study, we illustrated strong associations between the BMI and low-dimensional representations of functional connectivity, indicating plausible links between functional gradients and eating behaviors [31]. Recent advances in machine learning have made strides in feature representation to learn novel features from an existing set of features for various downstream machine-learning tasks. In particular, an autoencoder creates latent features that effectively describe the original features through nonlinear data compression and reconstruction [32–34]. Autoencoders have been adopted in some studies to distinguish populations of Alzheimer’s disease [35,36], schizophrenia [37,38], and autism [39] from healthy individuals. The feature representation capability of the autoencoder led to a higher performance in solving classification problems compared with conventional neuroimaging features.

In this study, we combined connectome manifolds with feature representation to identify subgroups of eating behavior traits. Briefly, we generated eigenvectors from the functional connectivity matrix and constructed an autoencoder model to identify subgroups with different behavioral traits. Subsequently, we compared the characteristics of eating behavior traits and degree of obesity among subgroups and assessed between-group differences in cortico-cortical and subcortico-cortical connectivity. Additionally, we assessed the reproducibility of our findings using an independent dataset.

## Result

We studied 424 healthy adults obtained from the enhanced Nathan Kline Institute-Rockland Sample database (mean ± standard deviation [SD] age = 47.07±18.89 yr; 67% female; mean ± SD BMI = 27.82±5.77 kg/m^2^, range 16.26–47.93 kg/m^2^) [40]. Details of the participant selection, image processing, and analysis are described in the *Methods* section. Reproducibility of the findings was validated using an independent dataset, Leipzig Study for Mind-Body-Emotion Interactions database, which contained 212 healthy adults (mean ± SD age = 38.97±19.80 yr; 35% female; mean ± SD BMI = 24.17±3.67 kg/m^2^, range 17.93–36.65 kg/m^2^) [41].

### Low-dimensional representation of functional connectivity and autoencoder-based feature representation

For each individual, we built a functional connectivity matrix by calculating the Pearson’s correlation of the time series between different brain regions defined using the Brainnetome atlas [42]. Considering 210 cortical parcels, we computed a low-dimensional representation of the functional connectivity (henceforth, eigenvector) [22] using dimensionality reduction techniques implemented in the BrainSpace toolbox (https://github.com/MICA-MNI/BrainSpace; see *Methods*) [43]. Individual eigenvectors were linearly aligned to the template manifold and computed using the group-averaged functional connectome [43,44]. We selected three eigenvectors (E1, E2, E3) that explained approximately 54% of the information of the template affinity matrix (**Figure 1A**). Similar to the previous findings based on the Human Connectome Project dataset [22,31,43], each eigenvector exhibited different cortical axes; the first, second, and third eigenvectors expanded from the primary sensory to association cortices (E1), from visual to somatomotor (E2), and from the multiple demand network to task-negative systems (E3), respectively.

**Figure 1.**
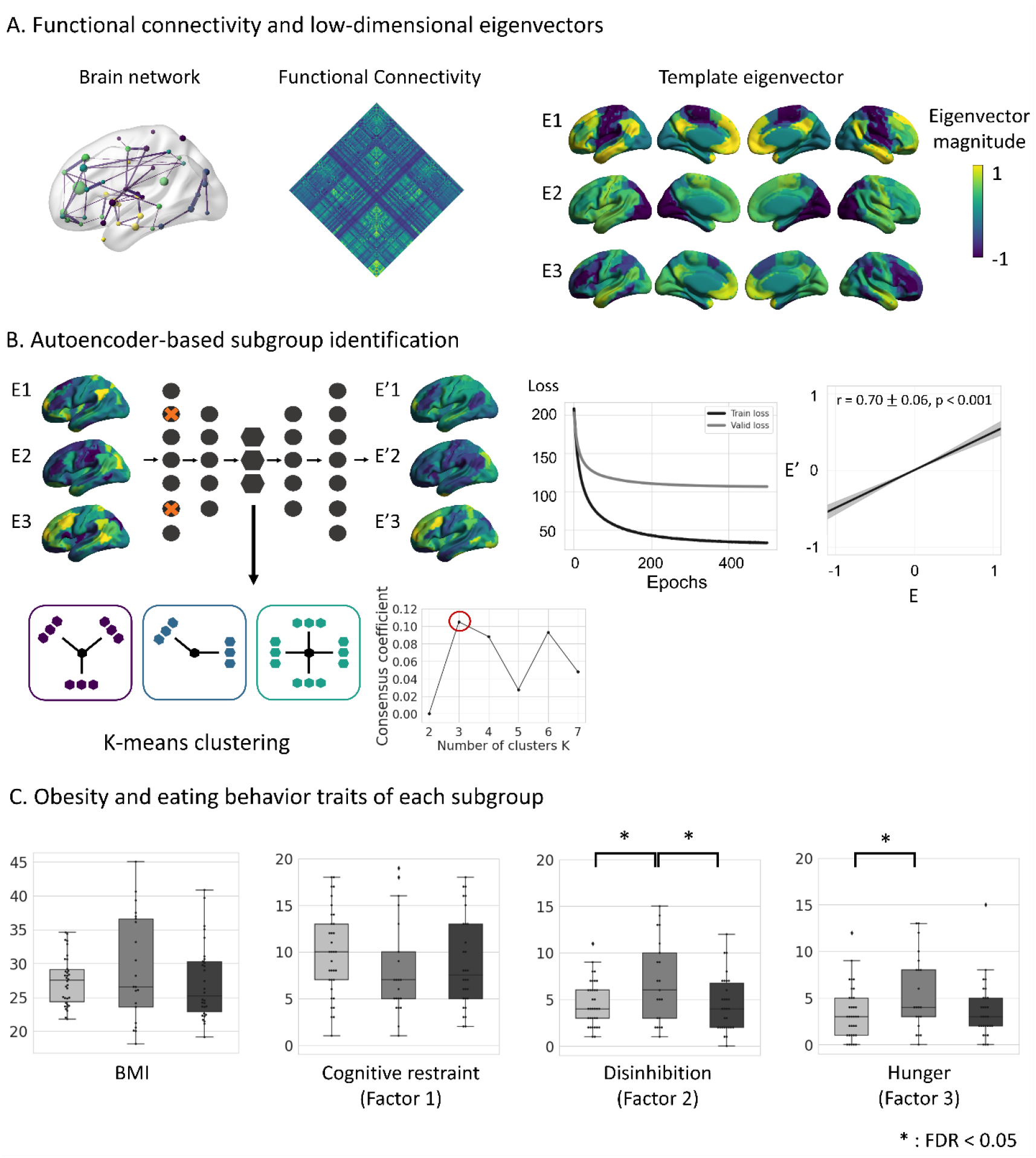
Subgroup identification using the manifold learning and autoencoder-based feature representation. **(A)** Schematic of functional connectome organization (left) and group averaged functional connectivity matrix (middle) are reported. Template eigenvectors were generated using dimensionality reduction techniques, and three dominant eigenvectors (E1, E2, E3) were selected. **(B)** The autoencoder model learned latent features of the eigenvectors after controlling for age and sex (left top), and loss values are plotted for each epoch (middle). We calculated linear correlations between the original (E) and reconstructed (E’) eigenvectors of the test dataset, and correlation coefficients across the subjects are reported with mean ± SD (right). We defined subgroups using the latent features of the autoencoder, where the number of clusters was determined using the consensus coefficient (left bottom). **(C)** Distribution of BMI and eating behavior scores of each subgroup is plotted. Significant differences in scores between subgroup pairs are indicated by asterisks. *Abbreviations*: BMI, body mass index; FDR, false discovery rate; SD, standard deviation.

The three generated eigenvectors were concatenated and used as inputs for the autoencoder model. The autoencoder model extracts latent features of the input through compression (i.e., encoding) and reconstruction (i.e., decoding) procedures (see *Methods*). The loss graph with respect to epochs demonstrated that the loss values decreased in both the training and validation datasets, indicating the appropriateness of model fitting. We applied the trained model at 499 epochs, which exhibited the highest performance in the validation dataset to the test data and found significant correlations between the original and reconstructed eigenvectors (mean±SD r = 0.70±0.06 across the subjects, p<0.001), indicating that the autoencoder learned the eigenvectors appropriately (**Figure 1B**).

### Subgroup identification using features from representation learning

The latent features learned from the autoencoder (i.e., features from the hidden bottleneck layer in the middle) were subjected to an unsupervised learning framework to identify subgroups of the study population. In particular, we employed the k-means clustering, and the number of clusters was determined using the consensus clustering approach, which was set to three [45] (see *Methods*; **Figure 1B**). To assess differences between the obesity-related traits across the identified subgroups, we compared the BMI and eating behavior scores of the subgroups based on a three-factor eating questionnaire (TFEQ) [46]. We found significant differences (false discovery rate (FDR)<0.05) in the disinhibition and hunger scales in one subgroup; however, BMI did not exhibit considerable differences (**Figure 1C**). Thus, the identified subgroups may reflect different eating behavior traits, independent of the degree of obesity.

### Interpretation of the latent features from representation learning

We utilized the integrated gradient interpretation model to explain the autoencoder-derived latent features [47]. The integrated gradient method computes the attribution of each element (i.e., the brain region) of the input to predict the output (i.e., latent features of the bottleneck) by progressively increasing the intensity of input values from a zero-information baseline to a particular intact input level and averaging attributions (**Figure 2A**) [47]. We used the integrated gradient technique to identify the brain regions that contributed to the latent features in the hidden layer that contained important information to reconstruct the original data (**Figure 2B**). We considered the three integrated gradient maps of eigenvectors and found that most higher-order networks, including limbic, dorsal attention, frontoparietal, and default mode networks, exhibited high contributions (**Figure 2C**). The results indicate that higher-order association and limbic regions greatly contributed to the reconstruction of the original eigenvectors.

**Figure 2.**
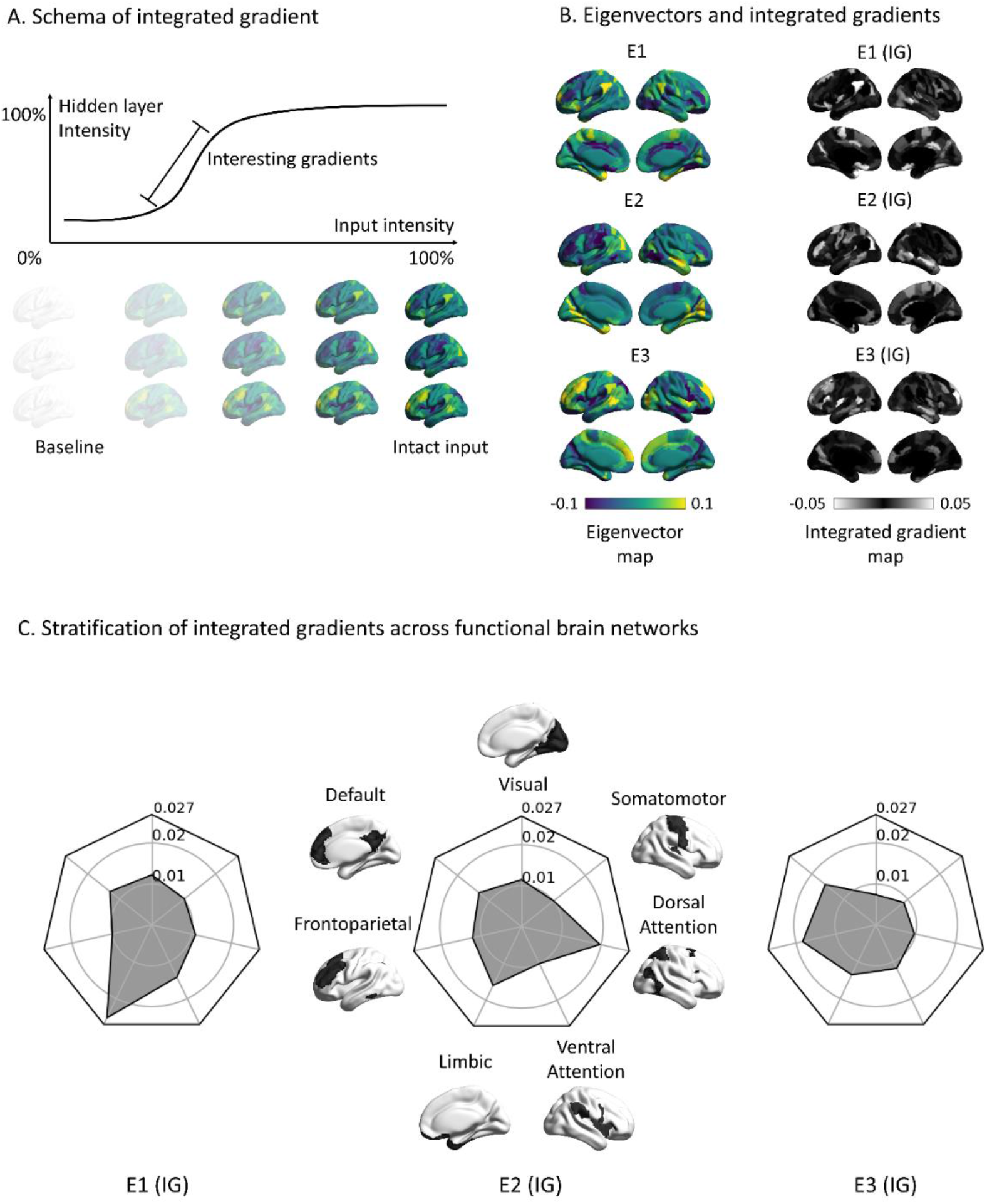
Characteristics of latent features using the integrated gradient technique. **(A)** Integrated gradient technique estimates the attribution of input towards predicting the output by averaging contributions while changing input intensities. **(B)** Spatial maps of each eigenvector and results of the integrated gradient technique are plotted on the brain surfaces. **(C)** Effects of integrated gradient are summarized according to functional communities. *Abbreviation*: IG, integrated gradient.

### Cortico-cortical and subcortico-cortical connectivity of subgroups

In addition to cortical alterations among subgroups, we hypothesized that connectivity in the reward circuit, which is known to be highly associated with the eating behavior may exhibit distinct profiles among subgroups. To prove our hypothesis, we investigated differences in the cortico-cortical connectivity based on integrated gradient maps among the three subgroups using the multivariate analysis of variance (MANOVA) [48]. We found significant differences in the precuneus, with the strongest and moderate effects in the frontoparietal and sensory regions (FDR<0.05; **Figure 3A**). Stratifying the effects according to the seven functional communities [49], somatomotor, ventral attention, frontoparietal, and default mode networks revealed strong between-group differences (**Figure 3A**). Additionally, we assessed between-group differences in the subcortico-cortical connectivity based on nodal degree centrality using ANOVA (see *Methods*). All subcortical structures exhibited considerable effects (FDR<0.05; **Figure 3B**), and stronger effects were observed in the accumbens, amygdala, and caudate (**Figure 3B**), which are involved in the reward system.

**Figure 3.**
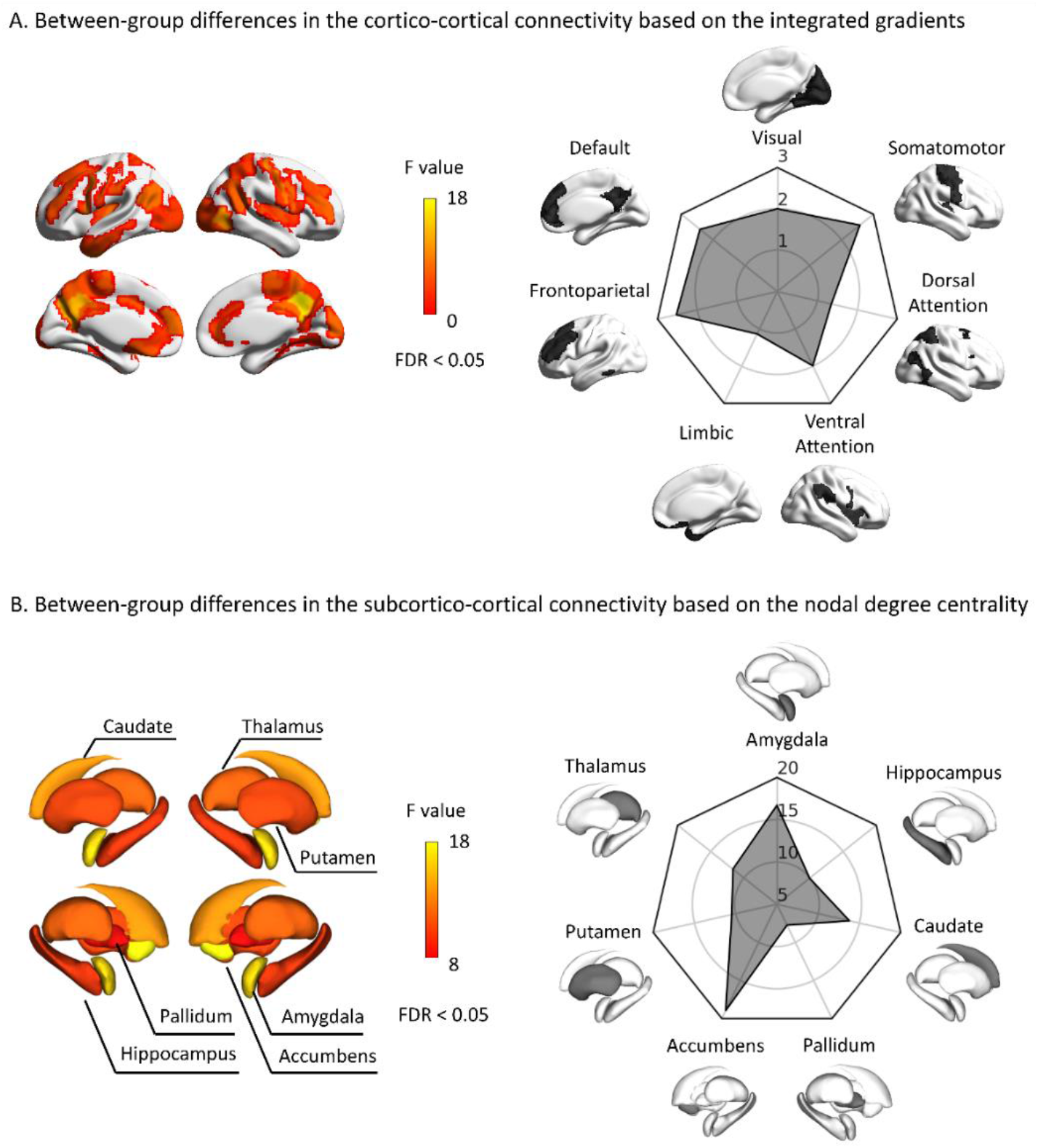
Between-group differences in the cortico-cortical and subcortico-cortical connectivity. **(A)** Between-group differences in the cortico-cortical connectivity based on the integrated gradients maps among the subgroups are visualized on the cortical surfaces, where the findings were multiple comparisons corrected using FDR<0.05. Effects were stratified according to seven intrinsic functional communities. **(B)** We visualized between-group differences in subcortico-cortical nodal connectivity strengths and stratified the effects according to each subcortical structure. *Abbreviation*: FDR, false-positive discovery rate.

### Cognitive associations

To provide the underlying cognitive associations of between-group differences in the cortico-cortical and subcortico-cortical connectivity, we utilized the Neurosynth meta-analysis cognitive decoding platform [50,51]. Associating the between-group differences in cortical and subcortical maps (**Figure 3**), we found high correlations with the reward-related terms, such as “anticipation,” “reward,” “incentive,” “monetary,” and “gain” (**Figure 4A**). Additionally, we associated the between-group difference maps with 24 cognitive state maps, as defined in [22], and observed strong correlations with “reward” (r = 0.40, FDR<0.05), and high associations with “emotion” (r = 0.24, FDR<0.05) and “affective” (r = 0.21; FDR<0.05; **Figure 4B**). Consequently, differences in the cortico-cortical and subcortico-cortical connectivity across the subgroups are associated with the reward-related cognitive functioning.

**Figure 4.**
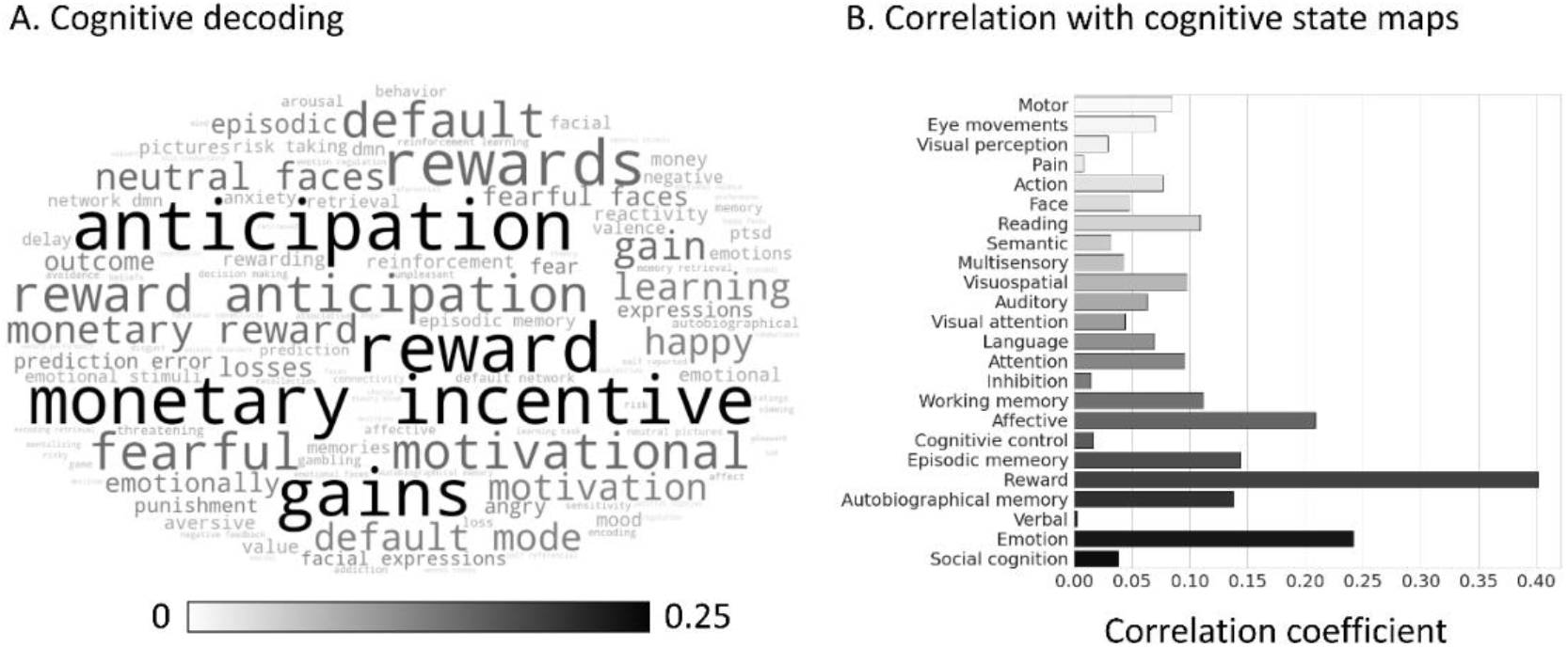
Cognitive associations. **(A)** We conducted cognitive decoding using the F-statistic map of cortico-cortical and subcortico-cortical connectivity differences across the subgroups using Neurosynth. **(B)** Correlation coefficients between the between-group difference maps and 24 different cognitive state maps are shown with bar plots.

### Replication of eating behavior traits

We performed the entire subgrouping analysis and compared the obesity and eating behavior traits across subgroups by using an independent dataset with different acquisition parameters (see *Methods*). One subgroup exhibited significant differences (FDR<0.05) in the hunger scale (**Figure 5A**), which particularly confirms the robustness of the hunger scale results. Notably, the subgroups defined from the two datasets exhibited comparable profiles (**Figure 5B**), with no statistical differences.

**Figure 5.**
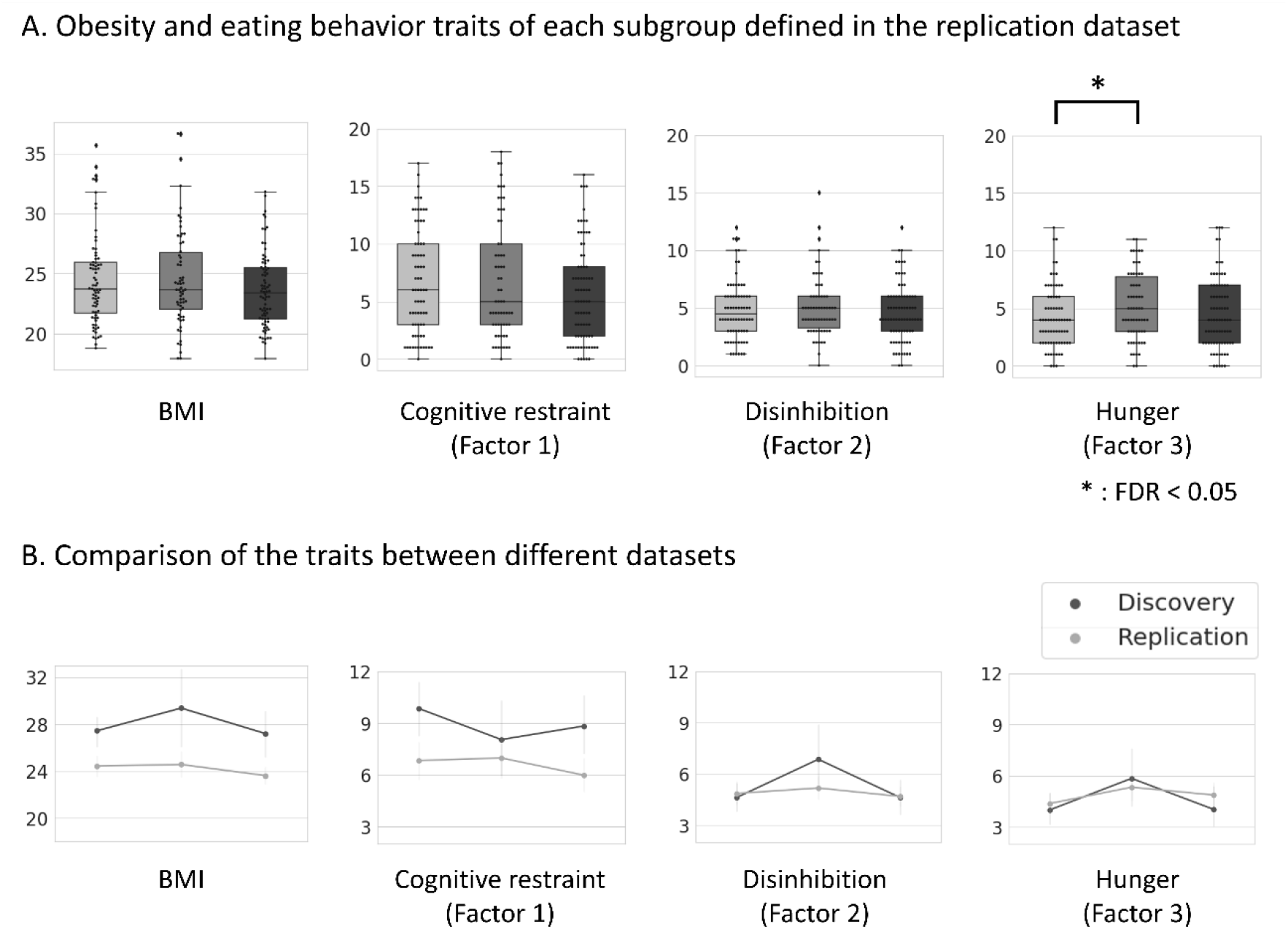
Reproducibility analyses. **(A)** Distribution of the BMI and eating behavior scores of each subgroup using the replication dataset are shown. Significant differences between the scores of subgroups are indicated by asterisks. **(B)** We compared profiles of the BMI and eating behavior scores of two different datasets. *Abbreviation*: BMI, body mass index; FDR, false discovery rate.

### Sensitivity analyses

Robustness of the findings was confirmed using the conducted analyses.

#### a) Subgroup identification without autoencoder

We applied k-means clustering to the concatenated eigenvectors and not to the latent features from the autoencoder. No significant differences were found between the BMI or eating behavior scores (**Figure S1**), indicating the necessity of applying the autoencoder model.

#### b) Bootstrapping analysis

We randomly selected 90% of participants with replacements and performed the same analyses of feature representation learning, clustering, and profiling of the obesity and eating behavior scores. We obtained consistent results that indicated the robustness of the findings (**Figure S2**).

#### c) Different densities of connectivity matrix

As in previous studies [22,26,43], our main findings were based on the functional connectivity with a density of 10%. Additionally, we performed the analyses based on the matrix with a density of 20 and 30% to evaluate the robustness and found consistent results (**Figure S3A**).

#### d) Different clustering methods

Instead of k-means clustering, we used the Gaussian mixture model clustering method, which is a probability distribution-based clustering approach (**Figure S3B**) and found that the obesity and eating behavior scores of each subgroup exhibited comparable profiles.

#### e) Different model architectures

We slightly changed the architecture of the autoencoder model and generated latent features. (i) We removed dropout layers; (ii) added one more layer; and (iii) removed one layer during the encoding and decoding processes (see *Methods*). The obesity and eating behavior scores were not considerably different and exhibited similar trends (**Figure S4**).

#### f) Manifold eccentricity

Rather than utilizing an autoencoder model to compress three eigenvectors into a single latent feature, we adopted the manifold eccentricity analysis, which depicts the distance of each brain region from the center of the template manifold [23,29] (**Figure S5A**). We defined subgroups based on the manifold eccentricity. Here, the BMI and eating behavior scores did not exhibit considerable differences, indicating that the latent features, which were defined using the autoencoder model were more useful for identifying behavioral differences across the identified subgroups (**Figure S5B**).

## Discussion

Eating behavior is highly associated with the brain function [10,31]; however, owing to the heterogeneity of eating behavior traits, no conclusions have been made to relate eating behaviors with the brain. In this study, an advanced technique combining the dimensionality reduction (i.e., connectome manifolds) with representation learning of the autoencoder was used to identify three subgroups with different eating behavioral traits independent of BMI. The latent features learned from the autoencoder were used to subdivide the groups, which yielded three distinct subgroups with different eating behaviors on the disinhibition and hunger scales. Furthermore, differences in the cortico-cortical connectivity from integrated gradients and subcortico-cortical nodal strengths among the identified subgroups were associated with reward-related cognitive terms, indicating that alterations in the association and reward systems may be related to the eating behavior heterogeneity. We demonstrated the reliability and generalizability of our findings using an independent dataset with different acquisition parameters and demographic characteristics.

Machine learning is a powerful tool for analyzing neuroimaging data, and manifold learning techniques are increasingly being used to describe macroscale functional connectome organization along the cortex [27–29,52]. We generated a series of low-dimensional eigenvectors, and the spatial patterns agreed with those of existing studies based on the Human Connectome Project data [22,31,52]. We extended previous studies by analyzing these eigenvectors using an autoencoder model consisting of encoding and decoding processes to generate latent features that contain highly compressed information of the input data. Using the latent features, we obtained three distinct subgroups, which exhibited considerable differences in eating behaviors. Indeed, our previous studies based on the conventional graph-theoretical approaches found that the functional connectivity of frontoparietal and executive control networks is associated with disinhibited eating behaviors and eating concerns [10,53]. In addition, we revealed that external stimulation of the dorsolateral prefrontal cortex affected the brain function in the frontoparietal network, which in turn yielded a reduced appetite [54]. Our current findings complemented previous studies in that eating behaviors are related to the function of higher-order brain regions. Additionally, we extended these studies by defining subgroups of study participants to investigate the heterogeneity of eating behavior traits. Interestingly, eating behaviors on the disinhibition and hunger scales exhibited significant between-group differences at a similar BMI. Therefore, although eating behavior is highly associated with obesity, the neurological underpinnings of eating behaviors may be different from those related to obesity. Further comprehensive investigations are needed to assess the convergence and divergence between obesity and eating behavior traits with respect to the brain function to explore the underlying neurological mechanisms of their relationships.

The interpretation of neural network models is important for accurate disease diagnosis and precise decision-making processes [57–59], which is uncertain because of the complex combinations of nonlinearities of the model [55,56]. Several attempts have been increasingly made [57,60–67]. Representative techniques include the layer-wise relevance propagation (LRP) and class activation map (CAM). LRP redistributes the relevance of each node from a particular layer to the previous layer using a top-down process [68], and CAM calculates the weighted sum of feature maps at the last convolutional layer, where the weights are calculated at a fully connected layer connected to the last convolutional layer using global average pooling [69]. These techniques have been used for the diagnosis of multiple sclerosis and Alzheimer’s disease [65,67]. However, LRP is sensitive to the choice of network architecture [47], and CAM requires global average pooling. To overcome this limitation, a recent study introduced gradient-weighted CAM (Grad-CAM) using gradients contributing to specific outputs as weights to provide an explanation [70]. Integrated gradients expanded the prior methods, enforcing a few axiomatic properties that have a invariance toward different neural network implementations [47]. In this study, we used the integrated gradients approach and observed that limbic, dorsal attention, frontoparietal, and default mode networks exhibited strong attributions, indicating that subgroups might present distinct functional organization in large-scale networks of higher-order association and limbic regions.

Additional cortico-cortical and subcortico-cortical connectivity analyses and cognitive decoding approved that association and reward networks exhibited significant between-group differences among the identified subgroups. These systems are related to eating behaviors [4,10,14,71–74], and thus, features attributed to these networks may provide benefits in identifying subgroups related to eating behaviors. In particular, reward systems are highly associated with eating behaviors, where an imbalance between food-related reward circuits and inhibitory control systems yields an increased sensitivity to food, leading to overeating and weight gain [12,75–87]. In addition, reward circuits are regulated by dopamine-related neurotransmitters, where the atypical organization of dopaminergic circuits in mesolimbic and association cortices was observed in individuals with obesity [4,88–94]. Similarly, serotonin-related neurotransmitters control eating behaviors by inhibiting the hanger-stimulating system [95,96]. These studies collectively suggest that reward-related cognitive systems could be target regions for regulating eating behaviors independent of the degree of obesity. However, expanded validation is required to explore the biological mechanisms associated with these macroscopic brain alterations.

In this study, we identified subgroups showing different eating behavior traits, regardless of the degree of obesity by using connectome manifolds and feature representation learning. The findings were robust for independent datasets, thus suggesting generalizability. Although we interpreted latent features derived from the autoencoder model based on an integrated gradient approach, this technique measures indirect contributions of the input data. More advanced techniques for the direct inference of the contribution of latent features need to be considered in future studies. Our results provide a new evidence for eating behavior-related macroscopic imaging signatures, independent of obesity.

## Method

### Participants

Imaging and phenotypic data were obtained from the enhanced Nathan Kline Institute-Rockland Sample database (NKI-RS) [40]. We excluded participants who did not provide complete demographic information, BMI, or TFEQ scores. Of the 650 participants, 424 were selected for this study. The proportion of individuals with healthy weight (18.5≤BMI<25 kg/m^2^), overweight (25≤BMI<30), and obese (BMI≥30 kg/m^2^) was 144:151:121. In addition, we obtained independent data from the Leipzig Study for Mind-Body-Emotion Interactions (LEMON) database [41]. Participants without complete demographic information or obesity-related scores were excluded. In this study, we used 212 out of 229 participants. Details are presented in **Table 1**.

**Table 1.** Demographic information of study participants.

| Categories |  | NKI-RS | LEMON | p value<sup>1</sup> |
| --- | --- | --- | --- | --- |
| Subject number |  | 424 | 212 |  |
| Sex<br>(Female:Male) |  | 282:142 | 75:137 | <0.001<sup>2</sup> |
| Age |  | 47.07±18.89<br>(18.15–85.62) | 38.97±19.80<br>(23–78) | <0.001 |
| BMI |  | 27.82±5.77<br>(16.26–47.93) | 24.17±3.67<br>(17.93–36.65) | <0.001 |
| TFEQ | Cognitive restraint<br>(Factor 1) | 8.67±4.94<br>(0 – 20) | 6.36±4.65<br>(0–18) | <0.001 |
|  | Disinhibition<br>(Factor 2) | 5.03±3.45<br>(0–15) | 4.82±2.61<br>(0–15) | 0.39 |
|  | Hunger<br>(Factor 3) | 4.26±3.36<br>(0–15) | 4.61±2.94<br>(0–12) | 0.17 |

Mean±SD with range (minimum–maximum) are reported.
<sup>1</sup>The p values are calculated based on two sample *t*-tests between the discovery and replication datasets.
<sup>2</sup>Chi-squared test.
*Abbreviations:* NKI-RS, enhanced Nathan Kline Institute-Rockland Sample; LEMON, Leipzig Study for Mind-Body-Emotion Interactions; BMI, body mass index; TFEQ, three-factor eating questionnaire.

### MRI acquisition

#### a) NKI-RS

All imaging data were obtained using a 3-T Siemens Magnetom Trio Tim scanner. The acquisition parameters of T1-weighted data were as follows: repetition time (TR), 1900 ms; echo time (TE) = 2.52 ms, flip angle, 9°; field of view (FOV), 250 mm × 250 mm; voxel resolution = 1 mm^3^ isotropic, and number of slices, 176. The rs-fMRI data were as follows: TR = 645 ms, TE = 30 ms, flip angle = 60°, FOV = 222 mm × 222 mm, voxel resolution = 3 mm^3^ isotropic, number of slices = 40, and number of volumes = 900.

#### b) LEMON

LEMON imaging data were acquired using a Siemens 3 Tesla scanner equipped with a 32-channel head coil. Scanning parameters of the T1-weighted data were as follows: TR= 5000 ms, TE = 2.92 ms, inversion time 1 (TI1) = 700 ms, TI2 = 2,500 ms, flip angle 1 (FA1) = 4°, FA2 = 5°, echo spacing = 6.9 ms, bandwidth = 240 Hz/pixel, FOV = 256 mm, voxel resolution = 1 mm^3^ isotropic, acceleration factor = 3, and number of slices = 176. The parameters of the rs-fMRI data were as follows: TR = 1400 ms, TE = 30 ms, flip angle = 69°, FOV = 202 mm, voxel resolution = 2.3 mm^3^ isotropic, number of slices = 64 slices, number of volumes = 657, and multiband acceleration factor = 4.

### Data preprocessing

#### a) NKI-RS

The T1-weighted and rs-fMRI data were preprocessed using the fusion of neuroimaging preprocessing (FuNP) volume-based pipeline, which combines the AFNI, FSL, and ANTs software [97–100]. The magnetic field inhomogeneity of the T1-weighted data was corrected, and nonbrain tissues were eliminated. The rs-fMRI data were preprocessed as follows: the first 10 s of the volume were discarded, and head movements were corrected. The FIX software was used to eliminate nuisance variables, such as the cerebrospinal fluid, white matter, head motion, and cardiac- and large-vein-related abnormalities [101]. The artifact-free rs-fMRI data were registered onto the preprocessed T1-weighted data and subsequently onto the MNI152 standard space. Spatial smoothing with a full-width-at-half-maximum of 5 mm was applied.

#### b) LEMON

The T1-weighted data preprocessing was performed based on Nipype; the details are described in (https://github.com/NeuroanatomyAndConnectivity/pipelines/tree/master/src/_lsd_lemon) [102]. In brief, CBS Tools [103] were used to remove the background from the T1-weighted image, and masked images were used to reconstruct cortical surfaces using FreeSurfer [104,105]. The T1-weighted data were registered onto the MNI152 standard space based on the diffeomorphic nonlinear registration using ANTs [100]. The de-identification process was performed using CBS Tools [103] by applying a brain mask to all anatomical scans. The rs-fMRI data were preprocessed using Nipype [102]. The pipeline included the following steps: the first five volumes were discarded to allow for signal equilibration and steady-state conditions [106]. Head motion and MRI-induced distortions were corrected [99]. The rigid-body transformation was applied to co-register the rs-fMRI data with the anatomical image [107]. Denoising was based on Nipype rapidart and aCompCor [108], and band-pass filtering in the frequency range of 0.01–0.1 Hz was applied. Standardization of mean centering and variance normalization was performed [109], and the preprocessed data were registered onto the MNI152 standard space [100].

### Eigenvector generation

We constructed a functional connectivity matrix from the preprocessed rs-fMRI data (**Figure 1A**) by calculating Pearson’s correlation of the time series between two different regions. Brain regions were defined using the Brainnetome atlas [42] and a cortico-cortical functional connectivity matrix with a size of 210 × 210. The correlation coefficient was Fisher’s r-to-z transformation [110]. We generated principal eigenvectors of functional connectivity using the BrainSpace toolbox (*https://github.com/MICA-MNI/BrainSpace*) [43]. The diffusion map embedding algorithm [111], which is robust to noise and computationally efficient was used to estimate eigenvectors from the functional connectivity matrix, leaving only the top 10% elements per row. Eigenvectors of each individual were aligned to group-level template eigenvectors defined based on a group-averaged functional connectome via Procrustes alignment (**Figure 1A**) [43,44]. The age and sex were controlled using eigenvectors.

### Architecture of the autoencoder model

An autoencoder was used to generate latent features from concatenated eigenvectors. The autoencoder model is defined as follows:

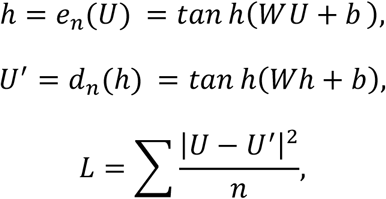

where *n* is the size of the input vector and *e_n_* is the encoder, which transforms the input vector *U* into a feature representation (or hidden representation bottleneck) layer *h*. Furthermore, *h* is used to reconstruct the input data and generate *U*’ using the decoder *d_n_*. The model was trained by minimizing the sum of mean square errors *L* between the input (*U*) and output (*U*’). The autoencoder model consists of two encoder layers, two decoder layers, and one feature representation layer. The feature representation layer had 210 latent variables, and each encoder and decoder layer had 630 and 420 units, respectively (**Figure 1B**). We used a hypertangent activation function in all layers, and a dropout rate of 0.3 was applied in the input layer [112]. The model was optimized using the Adam optimizer [113] with a learning rate of 1e^-4^, batch size of 10, and weight decay (i.e., L2-regularization) of 0.1. We concatenated three eigenvectors that provided sufficient information on the total functional connectivity data and entered it into the autoencoder model. We divided the dataset into training, validation, and test datasets with the ratios of 60, 20, and 20%, respectively. We trained the model using the training data and validated its performance using the validation data. We selected the model that exhibited the highest performance in the validation dataset for a total of 500 epochs. The selected model was applied to the test dataset, and its performance was assessed by calculating linear correlations between *U* and *U*’.

### Subgroup identification

Latent features in the hidden representation layer *h* were used to define the participant subgroups. We applied k-means clustering, which is based on the Euclidean distance. The optimal number of subgroups was determined using the consensus clustering, which robustly assesses how a pair of data is assigned to the same cluster (i.e., consensus coefficient) [45]. We determined the optimal number of subgroups, in which the largest consensus coefficient occurred, while varying the number of clusters. To assess the clinical and behavioral traits of subgroups, we compared the measured BMI and eating behavior scores using the TFEQ. The TFEQ consists of 51 questions [46], and each element is assigned to one of the three domains: (i) cognitive restraint, (ii) disinhibition, and (iii) hunger. We applied a two-sample *t*-test to compare each score between the subgroup pairs (**Figure 1C**). Significance was assessed using 1,000 permutation tests by randomly shuffling participants. A null distribution was constructed, and the real *t*-statistic value was deemed significant if it did not belong to 95% of the distribution (two-tailed p<0.05). Multiple comparisons were corrected using FDR [114].

### Integrated gradient for model explanation

To interpret latent features in the hidden representation layer, we assessed the attribution of each brain region to generate latent features using an integrated gradient approach (**Figure 2A**) [47]. The integrated gradient provides information on the extent to which a specific element (i.e., the brain region) in the input data contributes to predicting the output data (i.e., latent features). In particular, the integrated gradient (*IG*) from the *i*^th^ neuron is defined as follows:

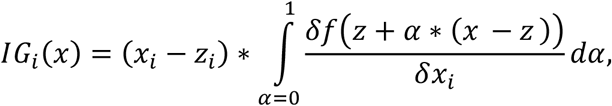

where *x* is the input data, *z* denotes the baseline, and *α* is the interpolation constant. The path integral can be approximated as follows:

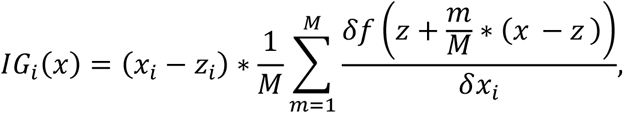

where *m* and *M* are the number of steps in the scaled feature perturbation constant and Riemann sum approximation of the integral, respectively. If the output has significantly changed, we assume that the attribution of the input data is high and vice versa. Thus, we can assess which brain regions of the input data considerably contributed during the compression and reconstruction processes. After calculating integrated gradients of each eigenvector, we stratified the effects according to the functional networks [49] to assess the networks that contributed the most to the reconstruction of the original data (**Figures 2B and C**).

### Between-group differences in cortico-cortical and subcortico-cortical connectivity

We compared the cortico-cortical connectivity computed from the three integrated gradient maps across subgroups using MANOVA (**Figure 3A**), where multiple comparisons were corrected using FDR [114]. The effects were stratified according to seven intrinsic functional networks [49]. In addition, we compared the subcortico-cortical connectivity across the subgroups using ANOVA (**Figure 3B**). The subcortico-cortical connectivity was quantified using the nodal degree centrality, a widely used graph-theoretical measure calculated by summing the connectivity strength of a particular brain area [115–117]. The nodal degree was estimated from the functional connectivity matrix, leaving only the top 10% elements per row. Multiple comparisons were corrected using FDR.

### Associations with cognitive states

Additionally, we assessed the relationships among the between-group differences in the cortico-cortical and subcortico-cortical connectivity across subgroups with cognitive terms using Neurosynth [50,51]. Neurosynth decodes the input data based on a meta-analytical method and provides correlation coefficients, whose cognitive terms are related to the data (**Figure 4A**). To systematically assess hierarchically organized cognitive maps, we performed spatial correlations between the between-group difference map in eigenvectors and 24 cognitive state maps, as defined in [22] (**Figure 4B**).

### Reproducibility experiments

We performed comparing eating behavior traits among subgroups to validate the generalizability of our results using the LEMON dataset [41]. We transferred the autoencoder model trained using the NKI-RS dataset into the LEMON dataset and applied the k-means clustering to identify subgroups. The obesity and eating behavior scores were profiled across the subgroups (**Figure 5A**), and the profiles were compared between the two datasets (**Figure 5B**).

### Sensitivity analyses

#### a) Subgroup identification without autoencoder

We applied the k-means clustering to the concatenated eigenvectors and not to latent features from the autoencoder in order to evaluate the effect of latent features on profiling clinical and behavioral traits (**Figure S1**).

#### b) Bootstrapping analysis

We performed 1,000 bootstraps with 90% resampled data to demonstrate the robustness of our results (**Figure S2**).

#### c) Different densities of connectivity matrix

We computed eigenvectors using different connectivity matrix densities of 20 to 30% and repeated the analyses (**Figure S3A**).

#### d) Different clustering methods

Instead of the k-means clustering, we used the Gaussian mixture model clustering approach, which creates clusters based on a probability distribution in order to assess the consistency of subgroup profiles (**Figure S3B**).

#### e) Different model architectures

We generated latent features by (i) removing the dropout layer (**Figure S4A**) and (ii) adding or (iii) subtracting one layer at the encoder and decoder (**Figures S4B, S4C**).

#### f) Manifold eccentricity

The same analyses were performed using the manifold eccentricity analysis [23,29], which computes the Euclidean distance between the center of the template manifold and all data points (i.e., brain regions) in the manifold space (**Figure S5**).

## Data availability

Imaging and phenotypic data were provided, in part, by the enhanced Nathan Kline Institute-Rockland Sample database, which is available after approval (http://fcon1000.projects.nitrc.org/indi/enhanced/index.html). Data from Leipzig Study for Mind-Body-Emotion Interactions database are publicly available at (https://ftp.gwdg.de/pub/misc/MPI-LeipzigMind-Brain-Body-LEMON/).

## Code availability

Codes for eigenvector generation are available in the BrainSpace toolbox (https://brainspace.readthedocs.io/en/latest/), surface visualization in the BrainNet Viewer toolbox (http://www.nitrc.org/projects/bnv/), and enigma toolbox (https://github.com/MICA-MNI/ENIGMA). Integrative codes for the full analyses are available at https://github.com/gudtls17/EatBehav.RepresentLearning.

## Author contributions

H. C., B. P., and H. P. designed the study, analyzed data, and wrote the manuscript. K. B. and J. L. aided in performing the experiments. S. H. reviewed the manuscript. B. P. and H. P. are the corresponding authors of this study and responsible for the integrity of data analysis.

## Supplementary Information

**Figure S1.**
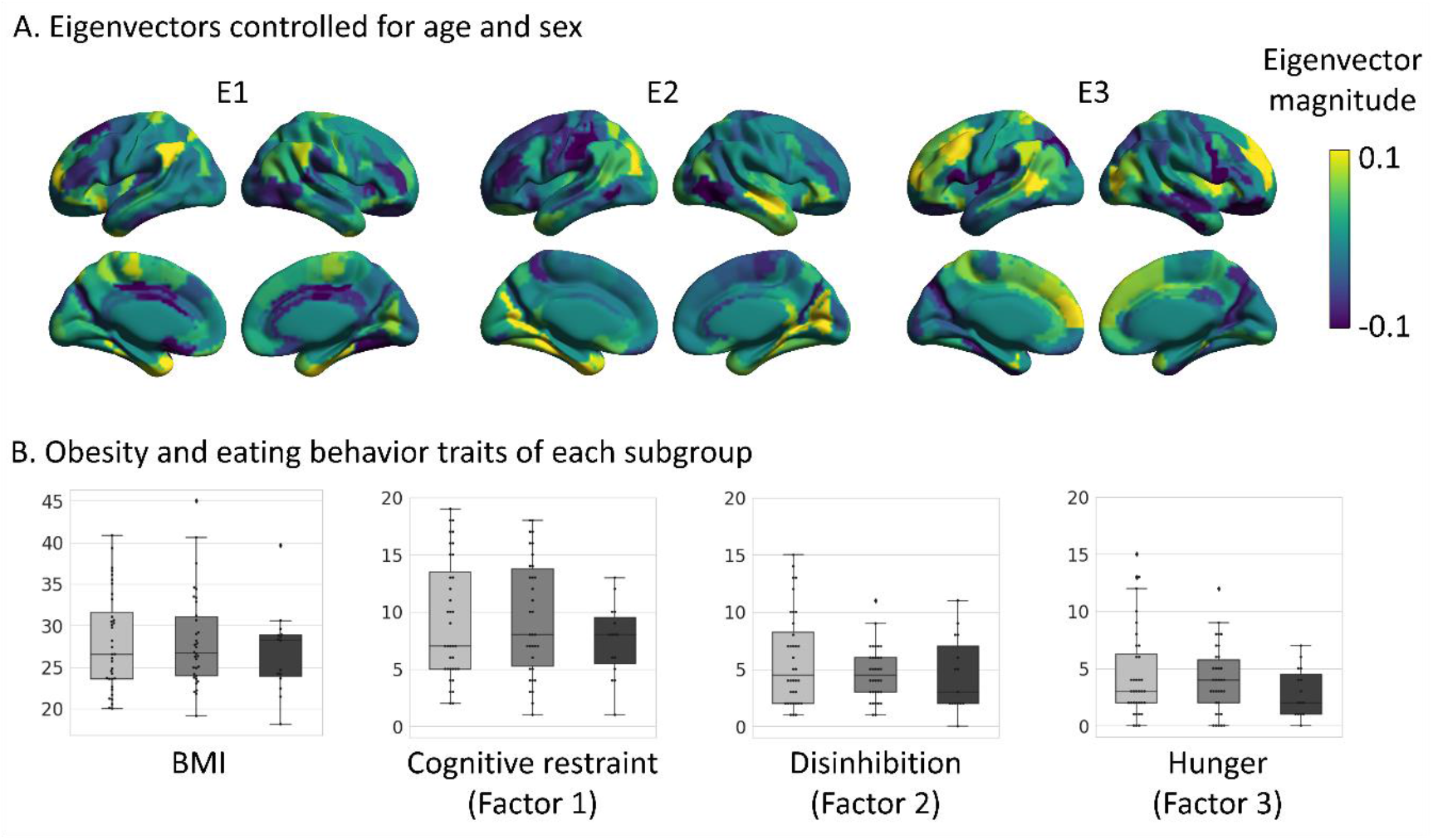
Subgroup identification without the autoencoder model. **(A)** Three eigenvectors (E1, E2, E3) controlled for age and sex are shown on brain surfaces. **(B)** Distribution of BMI and eating behavior scores of each subgroup are plotted. *Abbreviation*: BMI, body mass index.

**Figure S2.**
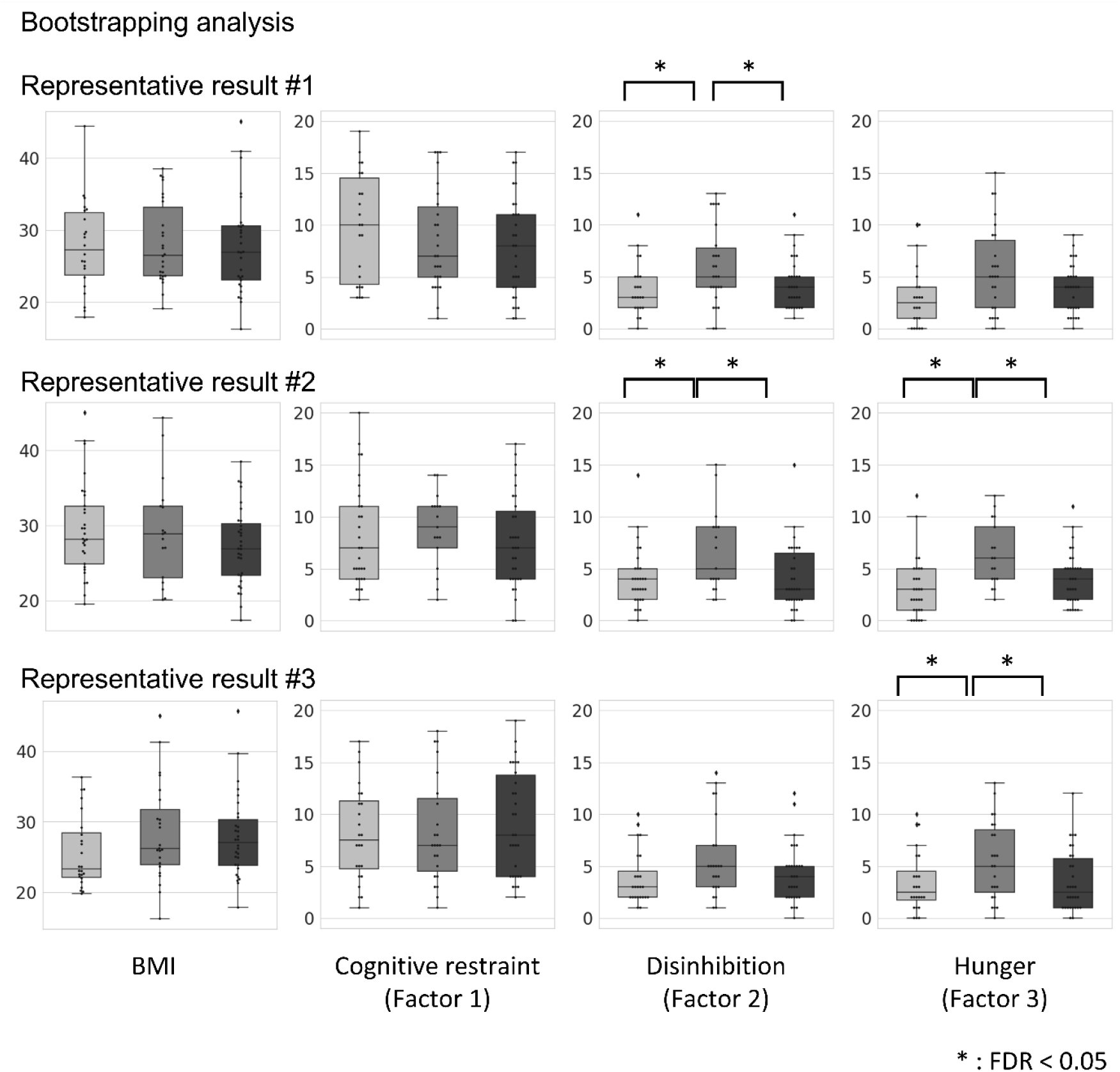
Bootstrapping analysis. We performed bootstrapping analysis by selecting 90% of participants with replacement and reported the obesity and eating behavior scores. Three representative results are presented. *Abbreviations*: BMI, body mass index; FDR, false discovery rate.

**Figure S3.**
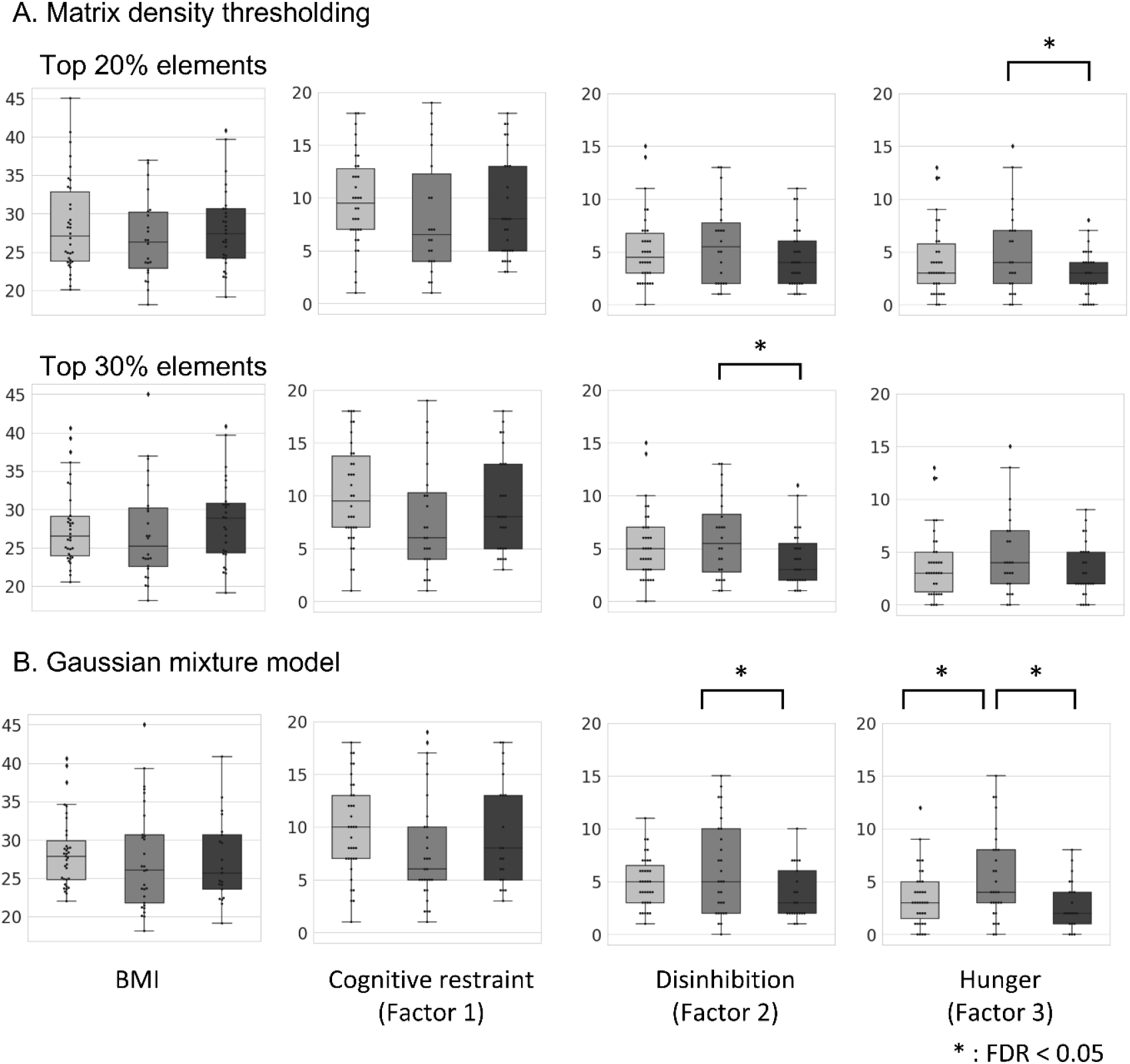
Different functional connectivity matrix densities and clustering method. We plotted BMI and eating behavior scores by changing the **(A)** matrix density with 20% (top) and 30% (bottom) and **(B)** clustering method to the Gaussian mixture model. *Abbreviations*: BMI, body mass index; FDR, false discovery rate.

**Figure S4.**
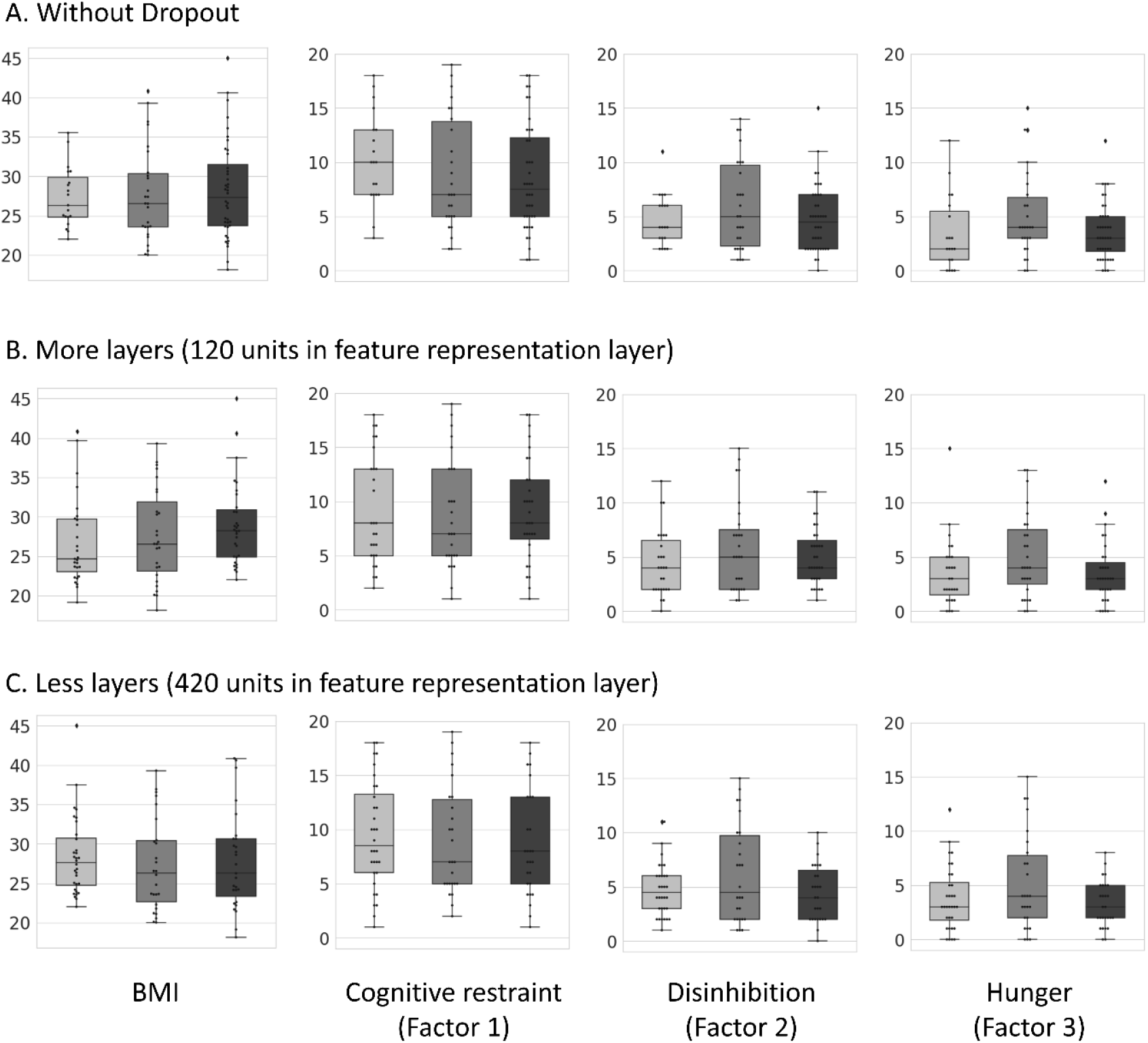
Different autoencoder models. **(A)** Distribution of the BMI and eating behavior scores without dropout layers. **(B)** Score distribution with one extra encoder and one extra decoder layers. The feature representation layer had 120 latent variables. **(C)** Score distribution with one less encoder and one less decoder layers. The feature representation layer had 420 latent variables. *Abbreviation*: BMI, body mass index.

**Figure S5.**
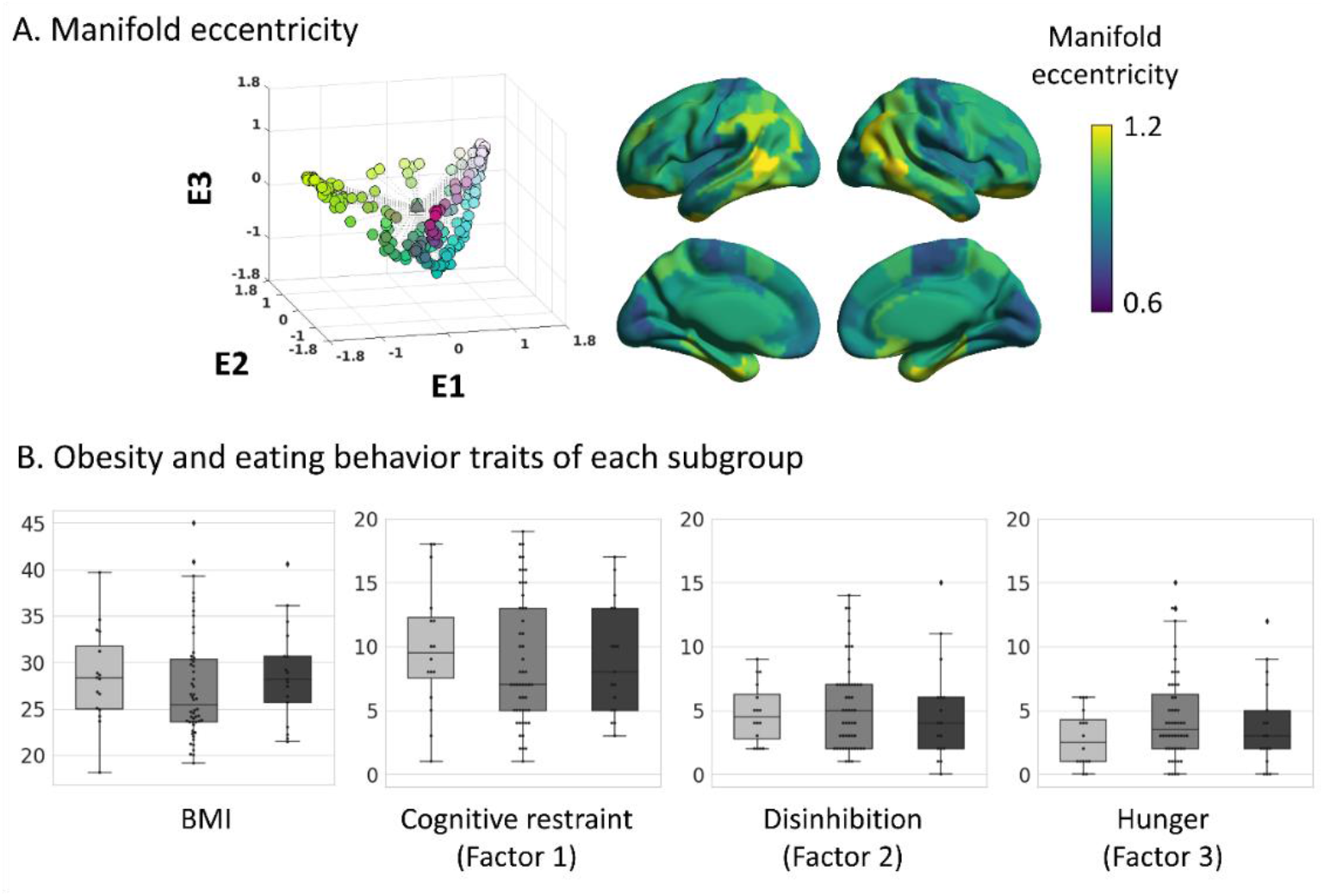
Subgroup analysis using the manifold eccentricity. **(A)** Dots in the scatter plot represents each brain region projected onto the three-dimensional manifold space, and colors are mapped onto the brain surface for visualization. **(B)** Distribution of the BMI and eating behavior scores are plotted. *Abbreviation*: BMI, body mass index.

